# Maternal RND3/RhoE deficiency impairs placental mitochondrial function in preeclampsia by modulating the PPARγ-UCP2 cascade

**DOI:** 10.1101/2020.06.22.164921

**Authors:** Liping Huang, Yanlin Ma, Lu Chen, Jiang Chang, Mei Zhong, Zhijian Wang, Ying Sun, Xia Chen, Fei Sun, Lu Xiao, Jianing Chen, Yingjun Lai, Chuming Yan, Xiaojing Yue

## Abstract

Preeclampsia (PE) is a life-threatening disease of pregnant women associated with severe hypertension, proteinuria, or multi-organ injuries. Mitochondrial-mediated placental oxidative stress plays a key role in the pathogenesis of PE. However, the underlying mechanism remains to be revealed. Here, we identify Rnd3, a small Rho GTPase, regulating placental mitochondrial reactive oxygen species (ROS). We showed that Rnd3 is down-regulated in primary trophoblasts isolated from PE patients. Loss of Rnd3 in trophoblasts resulted in excessive ROS generation, cell apoptosis, mitochondrial injury, and proton leakage from the respiratory chain. Moreover, Rnd3 overexpression partially rescues the mitochondrial defects and oxidative stress in human PE primary trophoblasts. Rnd3 physically interacts with the peroxisome proliferators-activated receptor γ (PPARγ) and promotes the PPARγ-mitochondrial uncoupling protein 2 (UCP2) cascade. Forced expression of PPARγ rescues deficiency of Rnd3-mediated mitochondrial dysfunction. We conclude that Rnd3 acts as a novel protective factor in placental mitochondria through PPARγ-UCP2 signaling and highlight that downregulation of Rnd3 is a potential factor involved in PE pathogenesis.

## Introduction

Preeclampsia (PE) is a pregnancy complication that occurs after the 20^th^ gestational week and is characterized by maternal hypertension, proteinuria, and multi-organ injuries. It affects 5-7% of pregnancies and is the leading cause of perinatal morbidity and mortality(1). Most of the cases with PE are associated with fetal intrauterine growth restriction (IUGR) and a high risk of maternal cardiovascular disease later in life(2). Although the mechanisms responsible for PE’s pathogenesis have not been entirely clarified, placental oxidative stress and poor vascularization are thought to be its leading cause(2, 3). Oxidative stress is generated by the imbalance of reactive oxygen species (ROS) and the endogenous oxidant scavenging system(4). Excessive ROS accumulation is identified in PE pathogenesis as the contributor to endothelial dysfunction and placental inflammation response(5-7). Although the shallow invasion of trophoblasts to decidua and subsequently failed spiral artery remodeling participate in placental ischemia and oxidative stress, accumulating evidence has shown that mitochondrial dysfunction during PE also can induce an increase in the level of ROS and cause placental oxidative stress(8-10). Abnormalities in mitochondria’ morphology, including swelling mass and disappearance of cristae, have been observed in trophoblasts of PE placenta(11). However, a limited number of studies have investigated the molecular machinery of the mitochondrial ROS during PE.

Peroxisome proliferators-activated receptor γ (PPARγ) is a ligand-inducible transcription factor that plays critical roles in mitochondrial biogenesis, dynamics, and metabolism(12, 13). Loss of PPARγ in a mouse model caused embryonic lethality with major placental defects(14) due to its critical roles in placental vasculature and the differentiation of complicated trophoblast lineages(15-17). Recent studies have revealed that the circulating agonists’ levels of PPARγ were significantly reduced in PE patients(14, 18). In contrast, administration of rosiglitazone, a selective PPARγ agonist, could largely ameliorate PE in a RUPP-induced PE rat model(19). Nevertheless, the regulatory mechanism of the downregulated PPARγ signaling in PE remains unknown.

Rnd3 is a small GTPase, also called RhoE, that exhibits multiple regulatory functions in carcinogenesis, cardiovascular diseases, and neural development(20-25). *Rnd3* has been proposed to be a PE candidate gene based on a population study. A specific *Rnd3* SNP (rs115015150) was associated with PE, and several quantitative cardiovascular risk traits(26). Several studies illustrated that Rnd3 was involved with PE and regulating the proliferation, migration, and invasion of trophoblast cells(27-29). In the present study, we identified decreases of Rnd3 expression in placentae and primary trophoblasts from PE patients and indicated a novel protective role of Rnd3 in PE via maintaining mitochondrial function. The possible molecular mechanism is that RND3 protein physically interacts with and stabilizes PPARγ. Transcriptionally regulated by PPARγ, uncoupling protein 2 (UCP2) was involved in mitochondrial membrane potential and ROS generation regulation. The downregulated PPARγ-UCP2 cascade was identified and could be reversed by Rnd3 overexpression and rescued mitochondrial dysfunction in human PE primary placental trophoblasts. The findings revealed a new function of Rnd3 in PE and provided new insights into PE pathogenesis.

## Methods

### Human placental tissues

Human placental tissues were obtained and used for the present study with written patient-informed consent and approval by the Ethics Committee of Southern Hospital, Southern Medical University. PE was defined according to the criteria of the American College of Obstetricians and Gynecologists (ACOG), which required blood pressure ≥ 140/90 mmHg on two occasions at least 4 h apart after 20 weeks of gestation with proteinuria ≥ 0.3 g/24h, or new onset hypertension without proteinuria but with other organ damage, such as thrombocytopenia, kidney or liver dysfunction and pulmonary edema. PE placental tissues were obtained from 24 patients with full term terminated pregnancies with average gestational ages of 36.1 ± 2.8 weeks. The healthy placental tissues were obtained from 34 healthy full term pregnancies with average gestational ages 37.0 ± 1.1 weeks. To ensure that the placental samples were sterile, all placentas were obtained via c-section deliveries consistently. The placental tissues were collected from the mid-section between the chorionic and maternal basal surfaces at four different positions 2 cm from central umbilical cord. Vessels were trimmed from the tissues. Fresh tissue samples were immediately frozen by liquid nitrogen or processed by paraformaldehyde fixation and frozen embedding. Women with a history of smoking and drinking, or diagnosed with multiple gestations, or other complications including chronic hypertension, diabetes, autoimmune and cardiovascular conditions were excluded from the present study. 91.2% PE patients received antihypertensive treatment with Nifedipine (0.5 mg/kg/day) after clinical diagnosis of PE.

### Electron microscopy analysis

Fresh placental tissue samples were fixed with a buffer containing 2.5% glutaraldehyde (diluted from 25% glutaraldehyde, G5882, Sigma-Aldrich, USA) and were subsequently processed and embedded in LX-112 medium. The ultrathin sections were stained with uranyl acetate and lead citrate, and then were examined in an H-7650 transmission electron microscope (HITACH, Japan).

### Isolation of primary human trophoblasts

The isolation of primary human trophoblasts was performed as described previously(30). Briefly, the fresh placental tissues were dissected into small pieces and were washed with PBS to remove blood cells. Digestion was performed at 37°C, with the media containing 0.125% trypsin (diluted from 0.25% Trypsin-EDTA, Gibco 25200072, USA), 0.03% DNase (D4263, Sigma-Aldrich, USA) and 1% Penicillin-Streptomycin. The digestion were neutralized by 5% fetal bovine serum (FBS). Following five digestion times, the floating cells were harvested by centrifugation at 1,200 x g for 15 min and re-suspended in DMEM medium. The trophoblasts were purified with 5-65% Percoll density gradients (P1644-500 ml, Sigma, USA). The cell surface biomarker cytokeratin 7 (CK7) was used to identify the trophoblasts. A total of 20 immunofluorescent staining images were acquired in different fields by fluorescence microscopy. CK7 positive cells and DAPI-labeled nuclei in each image were counted by the LAS V4.0 software. The trophoblasts’ purity was determined by the ratio of the number of CK7 positive cells over that of the total cells.

### Cell culture and hypoxia treatment

Human primary trophoblasts were cultured in RPMI-1640 medium containing 10% fetal bovine serum (FBS) and 1% Penicillin-Streptomycin. BeWo cells (ATCC CCL-98, USA) were cultured in F-12K medium containing 2 mM Glutamine, 10% FBS, and 1% Penicillin-Streptomycin. The cells were maintained at 37°C with 5% CO_2_. Hypoxic cell culture was performed in a hypoxic chamber (MIC-101, Billups-Rothenberg Inc, CA, USA) with 1% O_2_ for 16 h.

### Expression vectors and adenoviral expression vectors

The human Rnd3 cDNA subcloning was performed to generate the expression vectors GFP-RND3 and Myc-RND3 as described previously(22, 24). Human PPARγ cDNA was subcloned into GV141 backbone (GeneChem, Shanghai, China) to generate the expression vector Flag-PPARγ. The AdMaxTM system was used for the generation of recombinant adenovirus carrying human *Rnd3* cDNA. Briefly, CMV-EGFP-Rnd3 and viral backbone plasmid pBHG were co-transfected into HEK293 cells, and subsequently, the recombinant adenoviruses were harvested and amplified in HEK293 cells.

### Mitochondrial respiratory function measurement

Mitochondrial respirations were measured at 37°C in a Seahorse XF24 Extracellular Flux Analyzer (Agilent, USA). The XF Cell Mito Stress Test Kit (103015-100, Agilent, USA) and XF24 FluxPak mini (102342-100, Agilent, USA) cartridge were used to determine mitochondrial oxygen consumption rates (OCR) in viable cells. The mitochondrial assay medium consisted of XF Base Medium (102353-100, Agilent, USA), 10 mM glucose, 5 mM sodium pyruvate, and 2 mM L-glutamine at pH 7.4. The OCRs were measured by subsequent addition of 2 μM oligomycin, 1 μM FCCP, and 1 μM antimycin A/1 μM rotenone.

### Reverse transcription and quantitative PCR

The mRNA transcripts were quantified by quantitative PCR analysis, as described previously(22). Total RNA was prepared by TRIzol extraction. The forward and reverse PCR primers (5′ to 3′) were presented as follows: *RND3* (human): CCAGCCAGAAATTATCCAGCA/GAGAACCCGAAGTGTCCCA; *GAPDH* (human): GAGTCAACGGATTTGGTCGT/TTGATTTTGGAGGGATCTCG; *PPARγ* (human): TCCACATTACGAAGACATTCCA/CGACATTCAATTGCCATGAG; *UCP2* (human): TGGGTTCAAGGCCACAGATG/CCATTGTAGAGGCTTCGGGG; *PGC1α* (human): AGCACTTCGGTCATCCCAG/CAGTTTATCACTTTCATCTTCGC; *NRF1* (human): ATGGAGGAACACGGAGTGAC/TCATCAGCTGCTGTGGAGTT; *TFAM* (human): CCGAGGTGGTTTTCATCTGT/CCGCCCTATAAGCATCTTGA; *PPARα* (human): CTGTCTGCTCTGTGGACTCA/AGAACTATCCTCGCCGATGG; *SOD1* (human): TGAAGGTGTGGGGAAGCATT/GTCACATTGCCCAAGTCTCC; *SOD2* (human): TTTTGGGGTATCTGGGCTCC/TCAAAGGAACCAAAGTCACGT. *GAPDH* expression levels were used for qPCR normalization. The expression levels were determined by the 2^−ΔΔCt^ threshold cycle method.

### Luciferase assay

The luciferase reporter vector with the 1,000 bp promoter of the human *UCP2* gene was generated with a GV238 backbone (GeneChem, Shanghai, China). The luciferase assay was conducted, as described previously(22). Each measurement was repeated three times. All results were normalized according to the co-transfected *Renilla* luciferase enzyme activity (E1910, Promega).

### Western immunoblotting, immunoprecipitation, and ELISA

The protein samples for western blot analysis were prepared as described previously(22), and the immunoblotting densitometry was quantified by the ImageJ Software (NIH, USA). For immunoprecipitation, 293T cells were co-transfected with myc-Rnd3 and flag-PPARγ expression vectors. The cells were lysed in RIPA lysis buffer containing protease inhibitors. Cell lysates were incubated with protein A/G magnetic beads (88802, Thermo Fisher Scientific, USA) and either mouse IgG (5415S, Cell Signaling Technology, USA) or anti-Myc-Tag antibody at 4°C overnight. The beads were washed with lysis buffer for 4 times and boiled with 4× SDS loading buffer. The samples were analyzed by immunoblotting and identified with an anti-PPARγ primary antibody. We used the conformation-specific secondary antibody to recognize only the primary antibody but not the heavy and light chain of the antibody used for immunoprecipitation.

The antibodies used for the present study were from the following sources: anti-Cytokeratin 7 (ab9021, Abcam, USA); anti-PPARγ (2443S), anti-UCP2 (89326S), anti-Rnd3 (3664S), anti-Lamin B1 (12586S), anti-Myc-Tag (2276S), and mouse anti-rabbit IgG (Conformation Specific, 5127S) from Cell Signaling Technology, USA. Equal protein loading for immunoblotting was verified by the β-actin blot’s intensity (ab8226, Abcam, USA). Concentration of 8-Isoprostane was assessed by the 8-isoprostane ELISA Kit (516351, Cayman Chemical, USA).

### Fluorescence staining

For histological analysis, fresh placental tissues were embedded in the Tissue-Tek O.C.T. Compound (4583, Sakura, USA). Frozen sections were used for dihydroethidium (DHE) staining (D1168, Thermo Fisher Scientific, USA). Cellular JC-1 (T3168, Thermo Fisher Scientific, USA) staining and DHE staining were performed with viable BeWo cells, respectively. TUNEL staining (11684795910, Roche, Germany) was performed in fixed cells for apoptosis detection.

### Statistical analysis

The data were expressed as the mean ± standard deviation (SD). An unpaired, two-tailed Student’s *t*-test was used for two-group comparison. The one-way ANOVA, followed by the Student-Newman-Keuls method, was used for multiple-group comparison. The mRNA transcripts of Rnd3, PPARγ and UCP2 in human placentas were analyzed with F-test and Wilcoxon rank-sum test. All analyses were conducted using GraphPad Prism 8.0 and SAS 9.4. A value of *P*<0.05 was considered for a significant difference.

## Results

### The expression of Rnd3 is downregulated in human placentae with PE

To investigate the role of Rnd3 in PE, its expression levels were measured in 24 human placentae from PE patients and 34 healthy controls. Women with a history of smoking and drinking or diagnosed with chronic hypertension or gestational diabetes mellitus were excluded from the present study. The clinical characteristics of all pregnant women were presented in Table 1. Clinical data of blood pressure, BMI and proteinuria was collected at the time of PE diagnosis. In the placentae with PE, a 62.9% decrease was noted in the RND3 protein levels (Fig.1A), and a 46.0% decrease was reported in the *Rnd3* mRNA level (Fig. 1B). This clinical observation suggested that Rnd3 may act as a potential regulator in the pathogenesis of PE.

**Figure 1.**
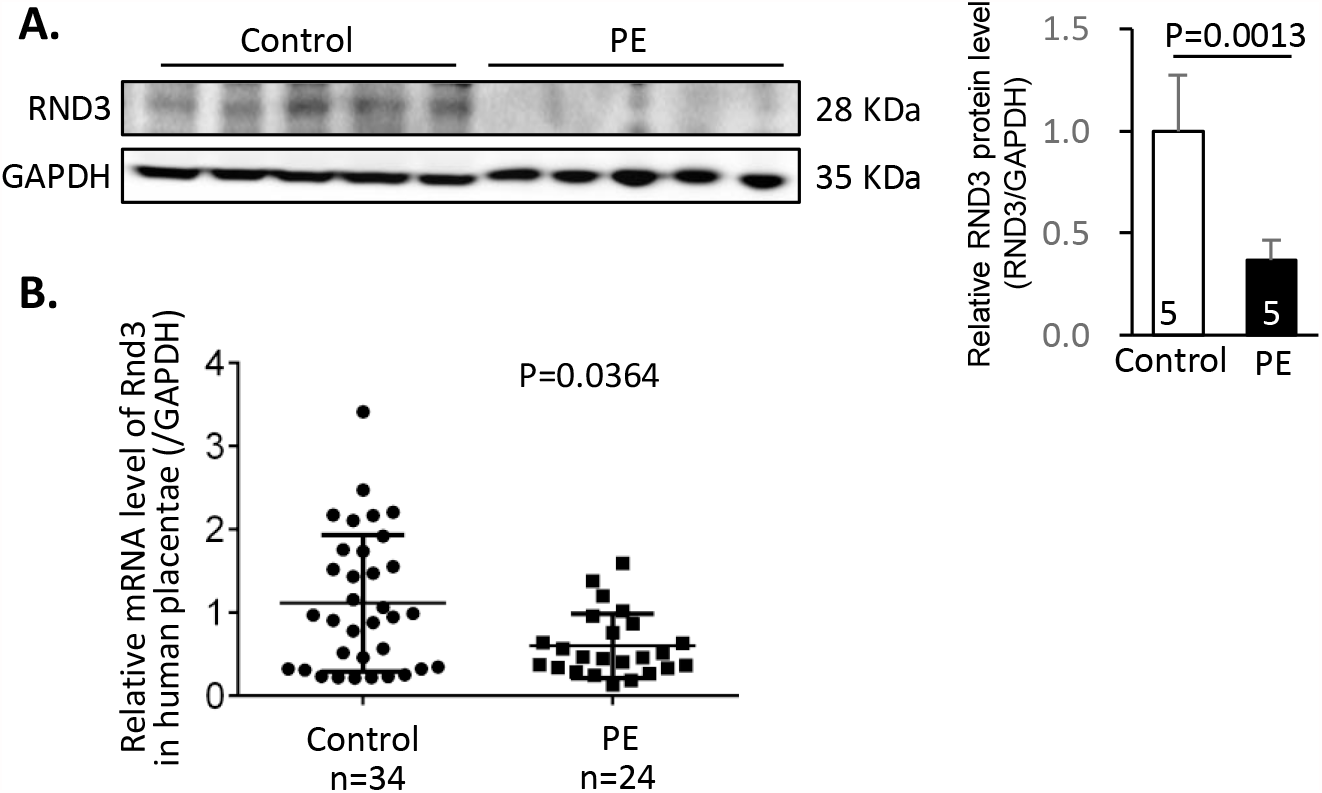
The expression of Rnd3 is downregulated in human placentae with PE. (**A**) Representative RND3 protein expression by Western blot analysis. Densitometry of bands was performed by Image J. (**B**) Relative levels of *Rnd3* mRNA from 34 human healthy placentae and 24 human placentae with PE.

### RND3 protein expression is downregulated in human PE primary trophoblasts and is associated with severe oxidative stress and apoptosis

Oxidative stress is the main regulator of PE development. Severe oxidative stress was noted in the placental tissue sections in the human placentae with PE (Fig. 2A). Systemic experiments based on primary human trophoblasts (PTBs) were conducted to understand the underlying mechanisms involved in this process. Human PTBs were isolated from PE and healthy placentas. The PTBs were identified by immunostaining the surface marker cytokeratin 7 (CK7) (Fig. 2B). According to nuclear counterstaining, the purity of the isolated PTBs was 97% (Fig. 2B). The protein level of RND3 in the PTBs was detected by immunostaining analysis. Rnd3 was universally expressed in the cytoplasm and nuclei of the trophoblasts (Fig 2C). Reduced Rnd3 expression was associated with increased ROS accumulation and was detected in the PE patient-derived PTBs (Fig 2C). To mimic the pathological PE placental environment, hypoxic cell culture of PTBs was established. Following hypoxia induction, additional ROS accumulation and apoptosis were observed in the PE PTBs compared with those noted in control PTBs (Fig 2D). This suggested that PE PTBs were more sensitive to hypoxic stress. Rnd3 may participate in the development of PE via the regulation of ROS generation.

**Figure 2.**
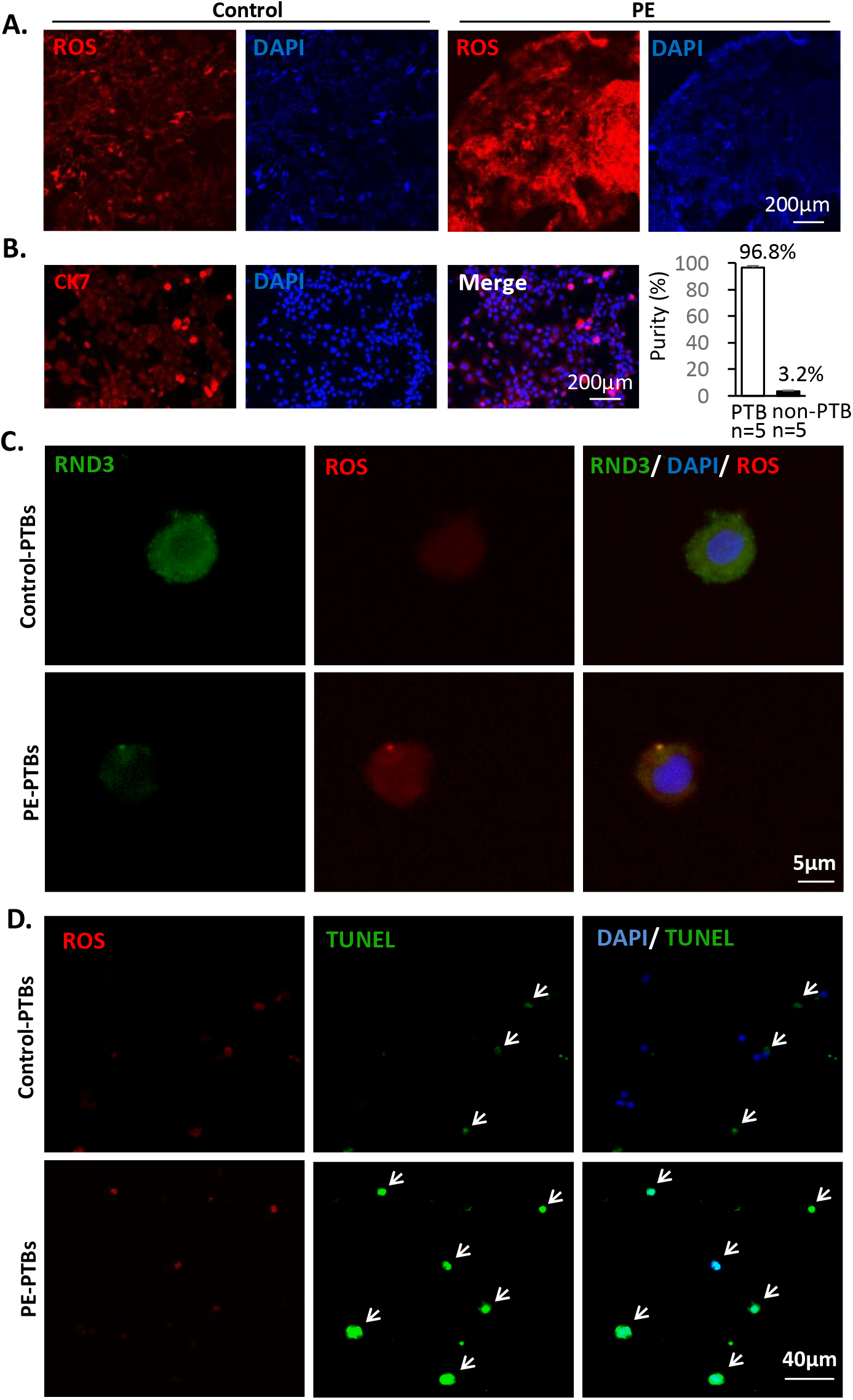
Decreased Rnd3 expression levels are associated with severe oxidative stress and apoptosis in PE patient derived-human primary trophoblasts. (**A**) Severe oxidative stress was observed in human placental tissue sections from PE patients by DHE staining. The nuclei were visualized by blue DAPI staining. Scale bar represents 200 *µ*m. (**B**) Isolated human primary trophoblast cells (PTBs) were identified by the cell surface marker CK7 shown in red and the nuclei were visualized by DAPI. The purity of primary trophoblasts was calculated by the ratio of CK7-positive cells over nuclei. The scale bar represents 200 *μ*m. (**C**) RND3 protein expression in PTBs was labeled by immunostaining. Decreased Rnd3 expression along with increased ROS levels were observed in the PTBs from PE patients. The scale bar represents 5 *μ*m. (**D**) Elevated ROS levels and severe apoptosis were observed in the PE PTBs compared with those of the PTBs from healthy control subjects. ROS was labeled by DHE staining shown in red. The arrows point at the TUNEL-positive cells (green), which overlapped with nuclear counter-staining (blue). The scale bar represents 40 *μ*m.

### Human PE primary placental trophoblasts are predisposed to mitochondrial damage

To explore the underlying mechanism of Rnd3-associated oxidative stress in PE, trophoblastic mitochondria, which are the main source of ROS, were analyzed by electron microscopy. In the placental tissues’ ultrathin sections, a borderline between the layers of syncytiotrophoblast (STB) and cytotrophoblast (CTB) could be identified. In the STB of the PE placentae, swelling mitochondria with loss of cristae (LOC) were observed, which were the typical characteristics of mitochondrial damage (arrows pointed, Fig 3A). We further analyzed the mitochondrial membrane potential in PTBs. Using JC1 staining, we compared the mitochondrial membrane potential between the control PTBs and the PE PTBs with or without hypoxic stimuli. The JC1 red/green fluorescence ratios between the two groups indicated no difference at the baseline (Fig 3B). The mitochondrial membrane potential was reduced in response to the hypoxic challenge. However, the red/green fluorescence emission ratio was collapsed in PE PTBs compared with that noted in control PTBs (Fig 3B), indicating that PE PTBs were predisposed to hypoxia-induced mitochondrial injury.

**Figure 3.**
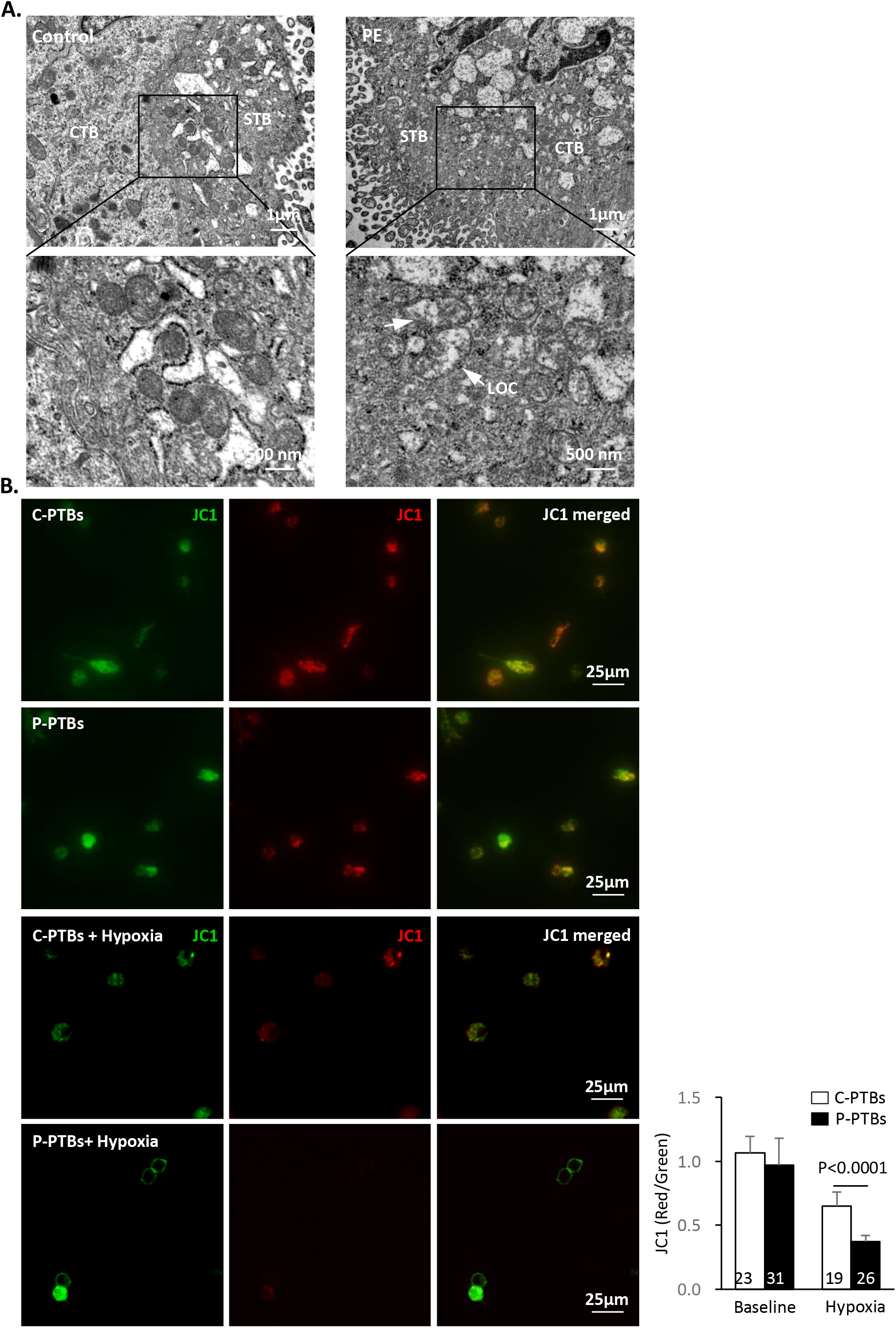
Abnormal mitochondria were observed in the human placental trophoblasts with PE. (**A**) Placental tissues from PE patients and healthy control subjects were analyzed by transmission electron microscopy and viewed at a low magnification (upper panel) and a high magnification (lower panel), respectively. The arrows point at the damaged mitochondria with loss of cristae (LOC). The scale bars represent 1 *μ*m and 500 nm, respectively. (**B**) Mitochondrial membrane potentials were shown by the JC1 staining of the primary trophoblast cells isolated from PE patients and healthy control subjects with or without hypoxic cell culture. JC1 is a cell permeable dye that accumulates in mitochondria and yields green fluorescence. Driven by high mitochondrial membrane potential, JC1 is able to enter the mitochondrial inner membrane and yield red fluorescence. The ratios of the red/green fluorescence intensities were quantified by the image J software. The numbers in the columns represent the numbers of cells in each group. C-PTBs indicates control-primary trophoblasts, and P-PTBs indicates PE-primary trophoblasts.

### Overexpression of Rnd3 protects the trophoblastic cells from ROS generation and induction of apoptosis

To investigate the potential role of Rnd3 deficiency in inducing oxidative stress in PE PTBs, we manipulated Rnd3 expression in the trophoblastic BeWo cell line. Hypoxic cell culture was applied to induce ROS generation. ROS levels were determined by DHE fluorescence labeling and the detection of cellular 8-isoprostane levels. The GFP-Rnd3 expressing BeWo cells displayed significantly decreased ROS levels than those of the surrounding non-GFP-Rnd3 cells (Fig 4A). Consistent with these observations, myc-Rnd3 overexpression led to decreased 8-isoprostane levels (Fig 4B) and ameliorated cell apoptosis (Fig 4C-D), indicating that Rnd3 modulated the production of ROS in trophoblasts. Rnd3 deficiency was observed in human PE PTBs and may cause severe oxidative stress and apoptosis in PE.

**Figure 4.**
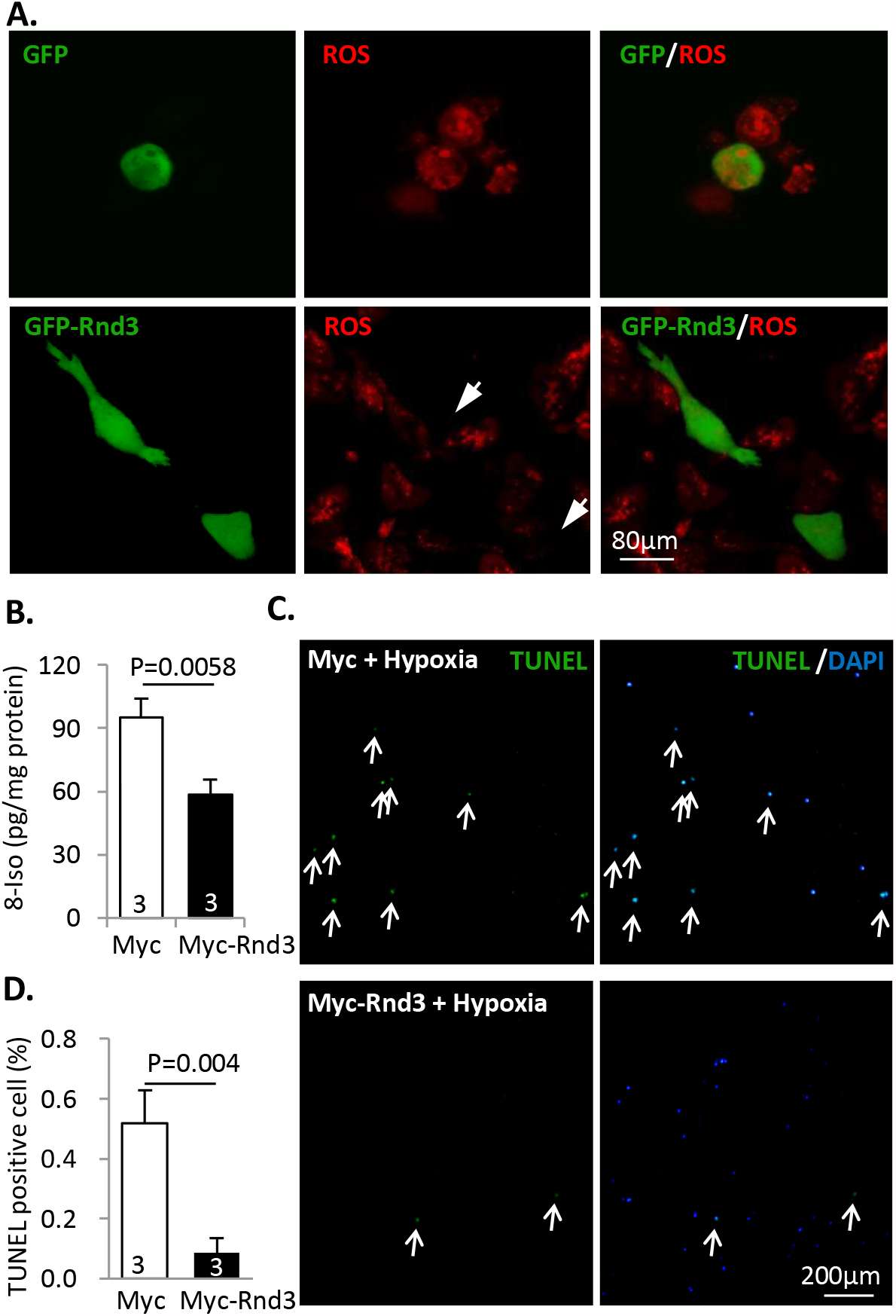
Rnd3 represses the hypoxia-induced ROS generation and apoptosis in trophoblastic BeWo cells. (**A**) Challenged with hypoxic condition, GFP-Rnd3 overexpressing BeWo cells exhibited reduced ROS levels. ROS was labeled by DHE staining shown in red. The arrows point at the GFP-Rnd3 expressing cells. The scale bar represents 80 *μ*m. (**B**) A decrease in 8-isoprostane levels was detected in the cell lysates of myc-Rnd3 overexpressing BeWo cells, compared with those noted in the myc control group. The experiments were repeated 3 times. (**C**) The comparison of the TUNEL staining in myc and myc-Rnd3 overexpressing BeWo cells following 16 h of hypoxic cell culture. The arrows indicate TUNEL-positive cells (green) overlapping with nuclear counter-staining. The scale bar represents 200 *μ*m. (**D**) Quantification of TUNEL-positive cells. The experiments were repeated 3 times.

### Rnd3 deficiency leads to mitochondrial damage in BeWo trophoblastic cells

To investigate if mitochondria contributed to Rnd3-mediated ROS generation, we knocked down Rnd3 in BeWo cells. Electron microscopy of Rnd3 knockdown BeWo cells revealed the mitochondrial injury. The compromised mitochondria displayed a structure with loss of cristae (LOC) and swelling mass (arrow pointed, Fig 5A). Consistent with the morphological changes, the mitochondrial transmembrane potential was also depolarized following Rnd3 knockdown (Fig 5B). Subsequently, the effects of Rnd3 on mitochondrial function were assessed by measuring the oxygen consumption rate (OCR) in different respiratory states. Comparing the Rnd3 knockdown group and the control group indicated no significant differences in the OCR of the basal respiration, maximal respiration, and space capacity (Fig 5C). However, a 2.4-fold increase in the proton leak associated OCR has detected in the Rnd3 knockdown BeWo cells (Fig 5C), indicating the uncoupling of ATP synthesis and substrate oxidation. Consistent with this result, the mitochondrial coupling efficiency was reduced in the siRnd3 group (Fig 5D).

**Figure 5.**
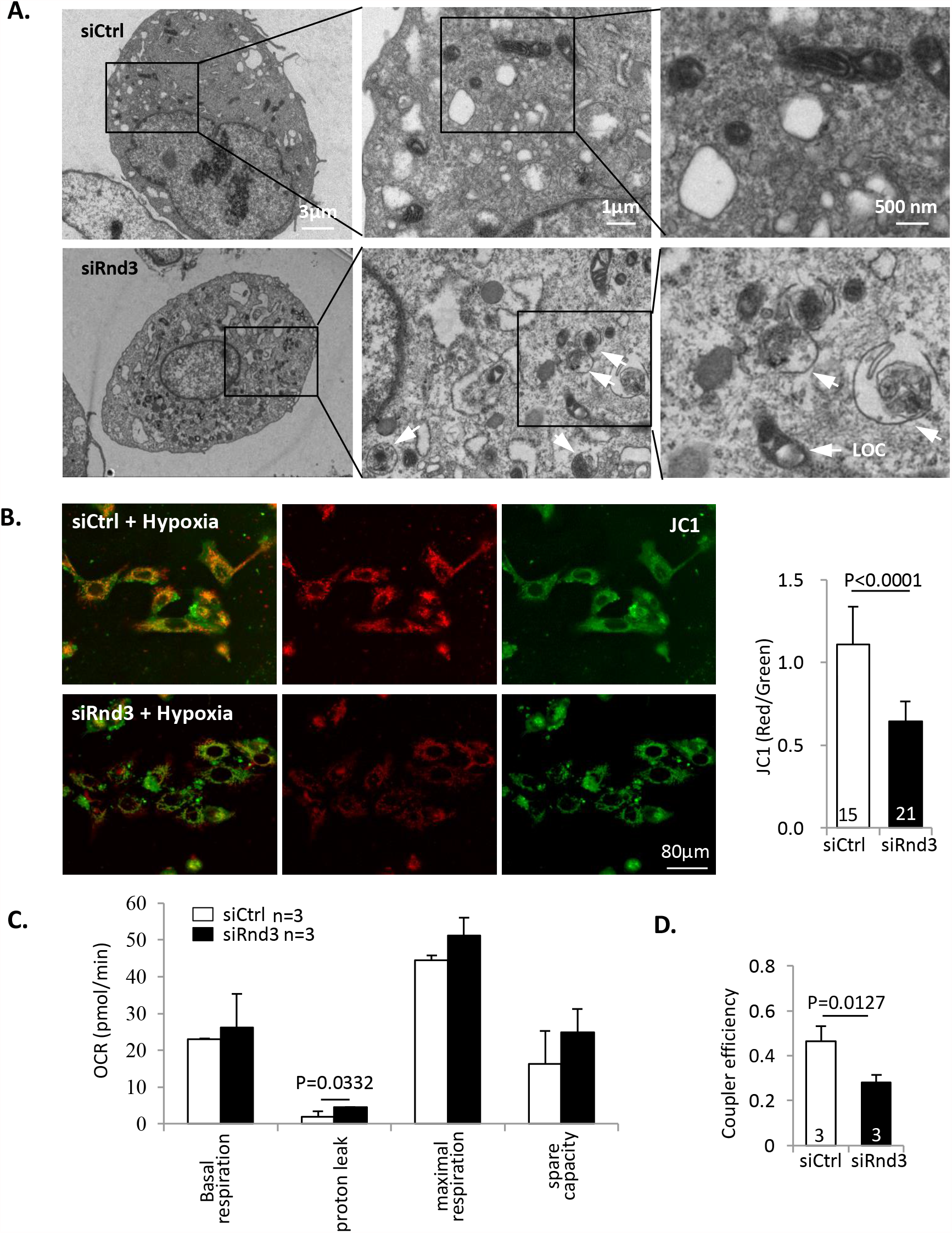
Rnd3 deficiency results in mitochondrial dysfunction. (**A**) By transmission electron microscope analysis, mitochondrial damage was observed in siRnd3 transiently transfected BeWo cells. The arrow heads indicate the loss of cristae (LOC) in the damaged mitochondria. The images of a series of magnifications are displayed. (**B**) The depolarized mitochondrial membrane potential was detected in siRnd3 transfected BeWo cells compared with that noted in the siCtrl group. The mitochondrial membrane potential was quantified by the ratios of red/green fluorescence intensities of mitochondria-specific JC1 dye. The scale bar represents 80 *μ*m. (**C**) The Oxygen consumption rate (OCR) of treated BeWo cells was measured prior to and following the injections of oligomycin, FCCP, rotenone and antimycin A (Rot/AA), respectively. Increased proton leak was detected in the siRnd3 group. (**D**) Reduced coupler efficiency was detected in siRnd3 transfected BeWo cells compared with that noted in the siCtrl group as determined by mitochondrial OCR determination. The numbers in the columns represent the number of cells in each group.

### Rnd3 facilitates the protein accumulation of PPARγ and stimulates the expression of UCP2 in trophoblasts

To explore the mechanisms of Rnd3-mediated mitochondrial dysfunction, the mRNA expression levels of the mitochondrial regulatory factors and ROS scavengers were evaluated in siRnd3 transiently transfected BeWo cells. The analysis indicated no significant difference between siRnd3 and control groups with regard to the mRNA levels of the transcription factors *PPARγ, PPARγ* coactivator 1-α (*PGC1-α*), peroxisome proliferator-activated receptor α (*PPARα*), nuclear respiratory factor 1 (*NRF1*), and mitochondrial transcription factor A (*TFAM*) (Fig S1). No significant differences were also noted about the levels of the endogenous ROS scavenger superoxide dismutase 1 (SOD1) and superoxide dismutase 2 (SOD2) (Fig S1).

Mitochondrial uncoupling protein 2 (UCP2) is critical for mitochondrial respiratory coupling. Rnd3 promoted the protein and the mRNA expression levels of *UCP2*, whereas knockdown of Rnd3 resulted in UCP2 deficiency (Fig 6A-6C), which was consistent with the mitochondrial respiratory coupling defect in siRnd3-transfected cells. As PPARγ has been identified as a transcriptional regulator of UCP2(31), we next assessed its expression levels. It is interesting to note that although the mRNA transcripts of PPARγ did not change the following manipulation of *Rnd3* (Fig S1), the PPARγ protein expression was stimulated by myc-Rnd3 overexpression and was repressed by Rnd3 knockdown (Fig 6A-6B), suggesting a possible post-translational regulation of PPARγ by Rnd3.

**Figure 6.**
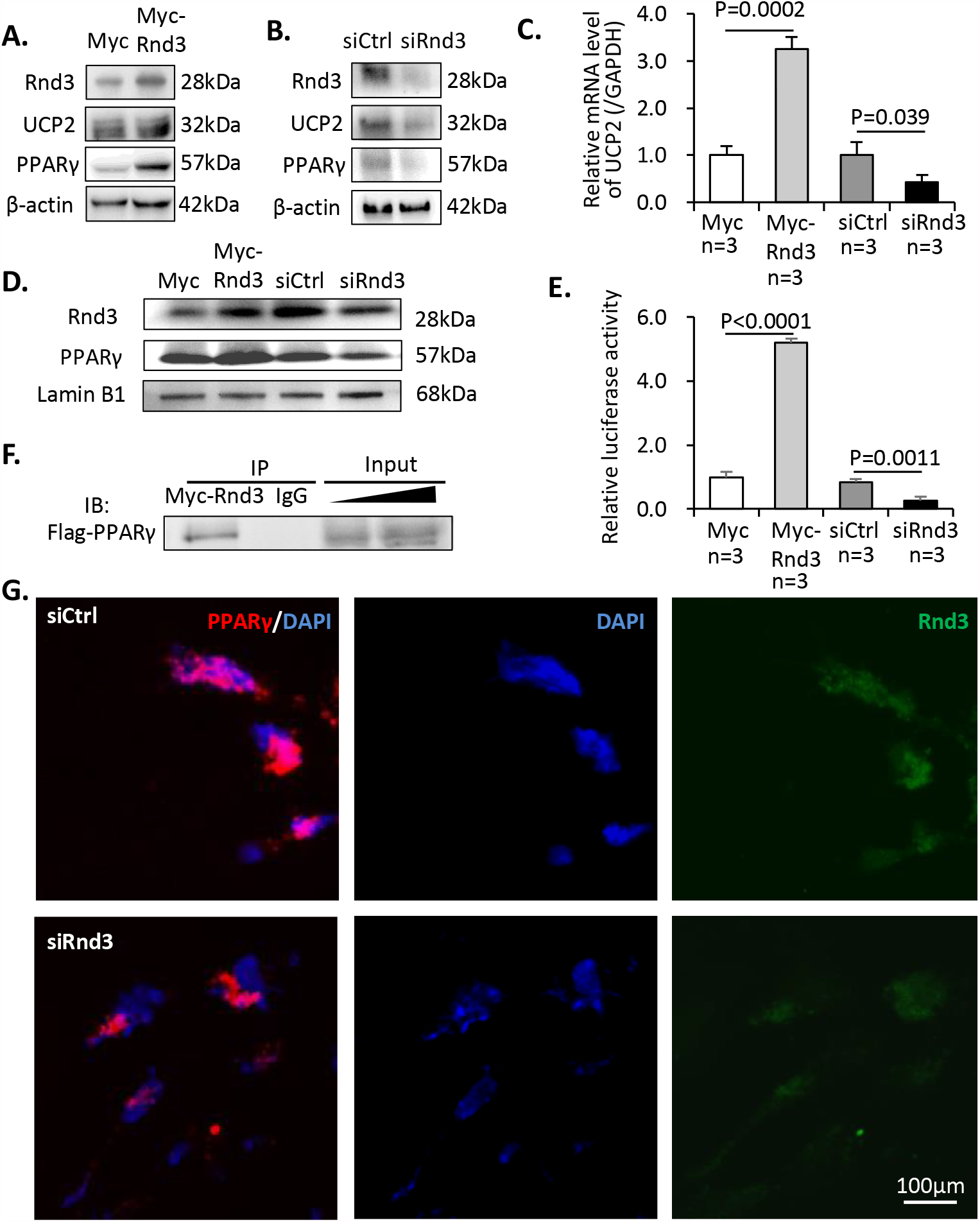
RND3 protein interacts with PPARγ and promotes the PPARγ-UCP2 cascade. (**A**) Increased expression levels of UCP2 and PPARγ were observed following Rnd3 overexpression. (**B**) Knockdown of Rnd3 results in lower UCP2 and PPARγ protein expression levels. (**C**) Q-PCR analysis indicated that *UCP2* mRNA levels were increased in myc-Rnd3 expressed cells, whereas they were reduced in Rnd3 deficient cells. (**D**) Nuclear protein levels of RND3 and PPARγ were increased in the myc-Rnd3 overexpressed cells and were decreased in the Rnd3 deficient cells. (**E**) Rnd3 regulated PPARγ transcriptional activity as demonstrated by luciferase assays. (**F**) Myc-RND3 physically interacted with flag-PPARγ *in vivo* as determined by coimmunoprecipitation. (**G**) Immunofluorescence staining indicated the co-localization of PPARγ and RND3 proteins. Knockdown of Rnd3 reduced the nuclear accumulation of the PPARγ protein. The scale bar represents 100 *μ*m.

### RND3 is physically bound to PPARγ and results in the accumulation of PPARγ in the nuclei

To investigate if Rnd3 could functionally regulate PPARγ protein, we knocked down and overexpressed the latter in BeWo cell cultures. As a transcription factor, the nuclear distribution of PPARγ is critical for maintaining its biological function. We detected apparent nuclear accumulation of PPARγ protein corresponding to the increase in Rnd3 expression. The opposite trend was observed when Rnd3 was knocked down (Fig 6D). Subsequently, a luciferase experiment was conducted to determine if Rnd3 caused any effect on its transcriptional activity. The luciferase reporter was driven by a *UCP2* promoter, which was responsible for the PPARγ-dependent transcriptional activity. Myc-Rnd3 overexpression resulted in a 5.2-fold increase in luciferase activity compared with that of the control, while downregulation of Rnd3 weakened PPARγ transcriptional activity and resulted in a 68% declined luciferase signal (Fig 6E). To further explore the potential mechanism of Rnd3-induced PPARγ protein accumulation, we performed an *in vivo* coimmunoprecipitation assay with 293T cells. As shown in Fig 6F, physical interaction between Myc-RND3 and Flag-PPARγ proteins was detected. Moreover, we performed co-immunostaining of these two proteins in BeWo cells and confirmed the co-localization of RND3 and PPARγ protein molecules and the reduced nuclear PPARγ protein levels in Rnd3 knockdown cells (Fig 6G).

### Mitochondrial dysfunction mediated by Rnd3 deficiency is attenuated by PPARγ overexpression

To assess whether PPARγ downregulation is responsible for Rnd3 deficiency-induced mitochondrial defects, Flag-PPARγ was overexpressed in Rnd3 knockdown BeWo cells. The improvement in mitochondrial membrane potential and mitochondrial morphological integrity was observed following PPARγ overexpression (Fig 7A-C). The ROS levels were detected by DHE labeling. We also detected 8-isopropane levels. PPARγ overexpression further ameliorated Rnd3 deficiency-mediated oxidative stress (Fig 7D-F). We analyzed the expression levels of UCP2 in Rnd3 knockdown BeWo cells with or without administration of Flag-PPARγ. As expected, PPARγ attenuated Rnd3 deficiency-induced UCP2 downregulation both at the mRNA and protein levels (Fig 7G-H). Finally, we assessed the mitochondrial respiratory functions of the different groups of BeWo cells. Overexpression of PPARγ significantly protected the mitochondrial function with improved proton leakage and coupler efficiency (Fig 7I-J).

**Figure 7.**
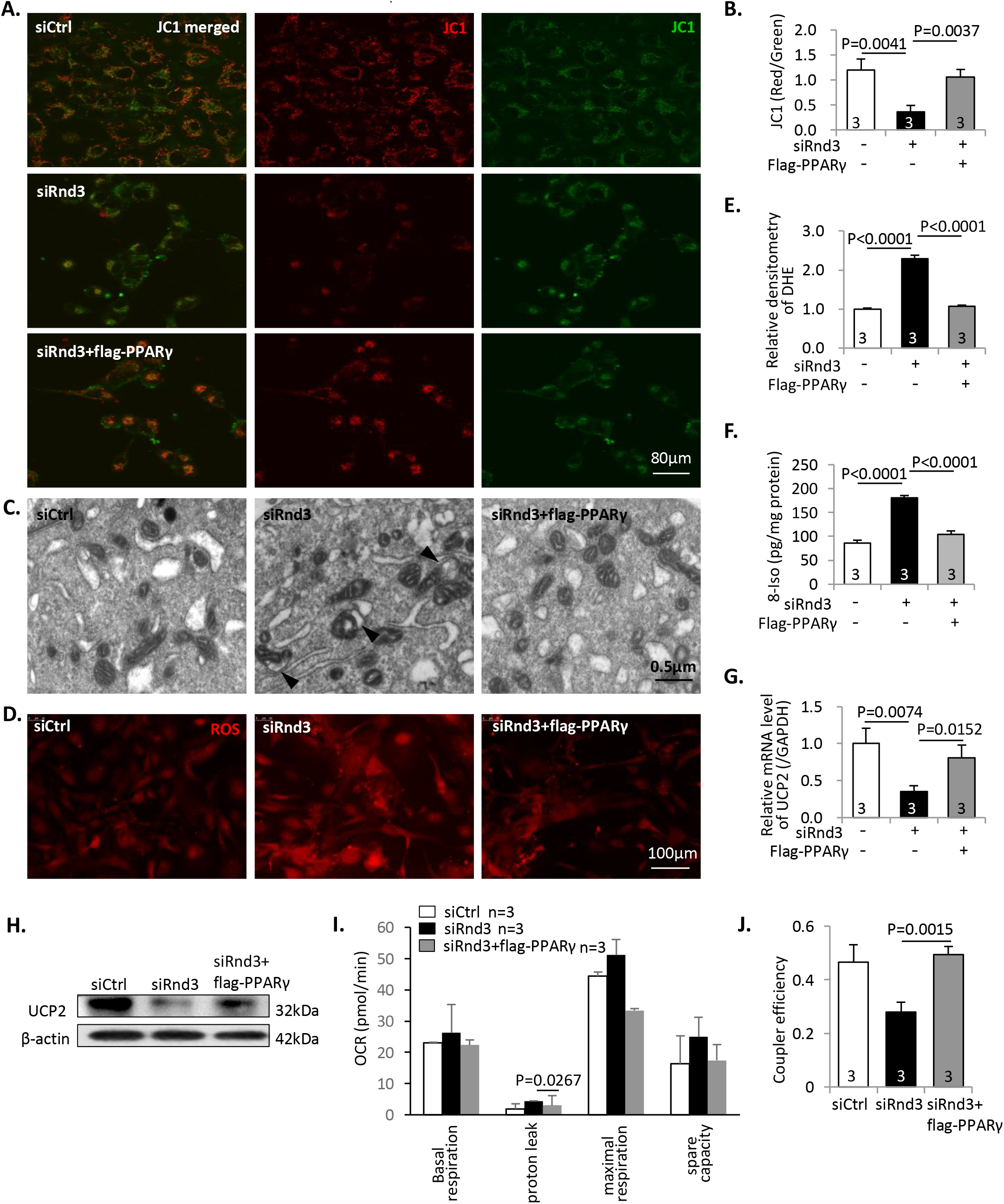
Rnd3 deficiency-mediated oxidative stress and mitochondrial dysfunction are attenuated by PPARγ overexpression. (**A**) JC1 staining was performed in BeWo cells among the three following groups: siCtrl, siRnd3 and siRnd3 plus flag-PPARγ. The depolarization of the mitochondrial membrane potential caused by Rnd3 knockdown was attenuated by flag-PPARγ overexpression. The scale bar represents 80 *μ*m. (**B**) The quantification of the ratios of red/green fluorescence intensities. The experiments were repeated 3 times. (**C**) Transmission electron microscope analysis indicates that the Rnd3 deficiency-induced mitochondrial damage was recovered by flag-PPARγ overexpression. The arrow heads display damaged mitochondrial structure in the siRnd3 group. The scale bar represents 0.5 *μ*m. (**D**) DHE staining indicates ROS levels among the three groups. The scale bar represents 100 *μ*m. (**E**) The ROS levels were quantified by the densitometry of DHE staining. The experiments were repeated 3 times. (**F**) 8-isoprostane levels were detected in the cell lysates of the three groups. (**G**) Q-PCR analysis of *UCP2* mRNA levels among the three groups. The experiments were repeated 3 times. (**H**) Immunoblots revealed increased UCP2 protein expression levels in the flag-PPARγ rescue group. (**I**) Mitochondrial respiratory function was measured. The proton leak associated OCR was improved by PPARγ. (**J**) Rnd3 deficiency-mediated defective coupler efficiency was ameliorated by PPARγ.

### Adenoviral-mediated Rnd3 overexpression in human PE primary trophoblasts rescues oxidative stress and mitochondrial defect

Rnd3 downregulation was demonstrated in human PE-PTBs along with severe oxidative stress and mitochondrial injury. To further reveal the critical role of Rnd3 in the clinical pathology of PE, we overexpressed hRnd3 in human PTBs by adenoviral-mediated gene delivery. Elevated RND3 protein levels in Ad-GFP-hRnd3 treated PTBs were confirmed by western blot analysis (Fig 8D). In the absence of Ad-GFP-hRnd3 infection, the baseline PE-PTBs exhibited higher ROS levels compared with those of the control-PTBs (Fig 8A, cells displaying no GFP, pointed by long arrows). Following infection with Ad-GFP-hRnd3, the ROS levels of the PE-PTBs were significantly reduced (Fig 8A, cells expressing GFP, indicated by arrowheads). The levels of 8-isopropane were decreased in PE-PTBs following Ad-GFP-hRnd3 application (Fig 8B), which was consistent with the previous observations. Increased expression levels of PPARγ and UCP2 were observed, as expected, in the PTBs with Ad-GFP-hRnd3 (Fig 8C-D). In addition to regulating the PPARγ/UCP2 pathway, Rnd3 further attenuated the mitochondrial function in PE-PTBs (Fig 8E-G).

**Figure 8.**
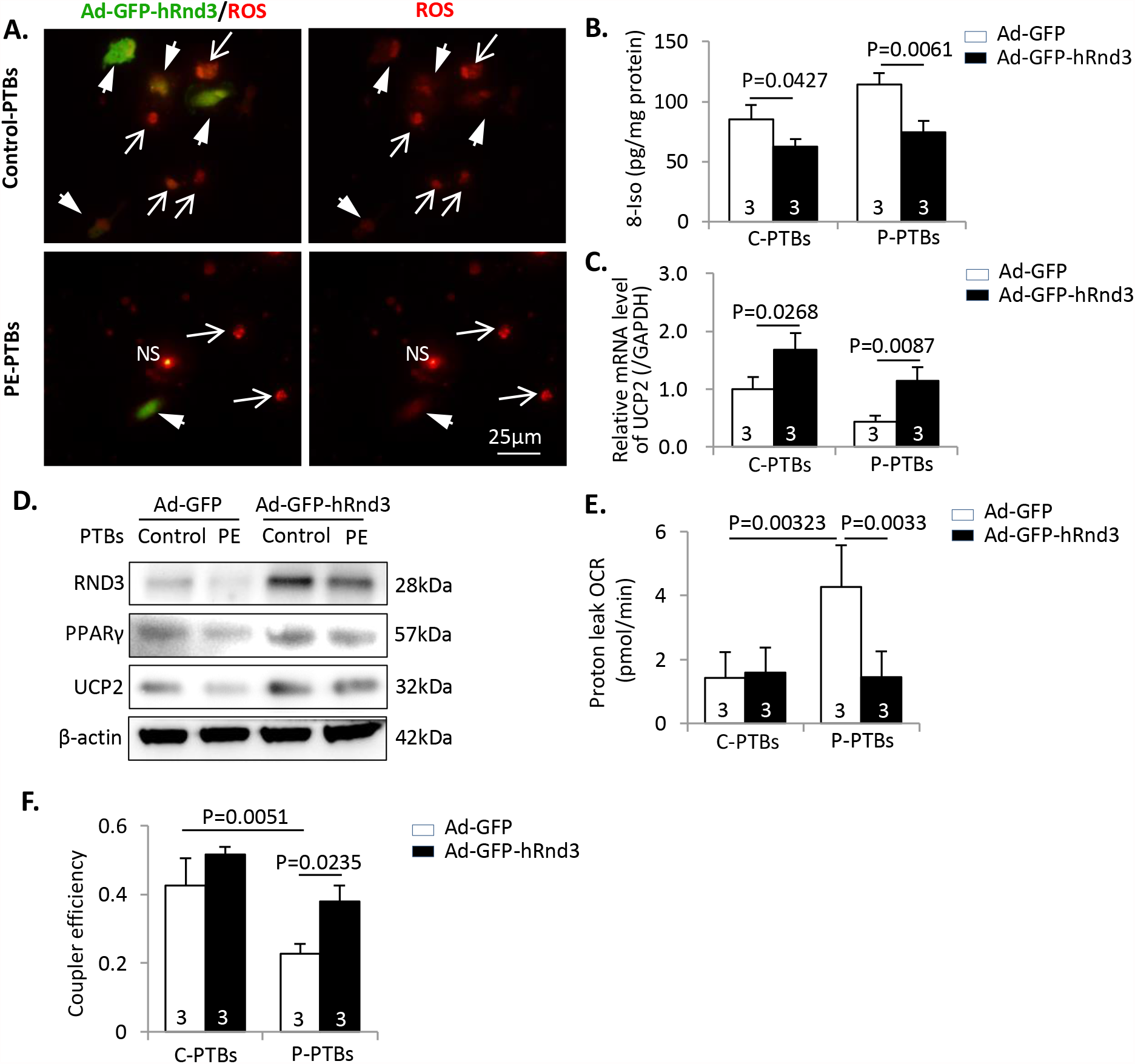
Oxidative stress and mitochondrial dysfunction in PE patient-derived PTBs are partially rescued by Rnd3 overexpression. **(A)** Human PTBs from PE patient and healthy control subjects were infected with Ad-GFP-hRnd3. Hypoxic cell culture was applied to induce oxidative stress. The arrow heads represent the GFP-Rnd3 expressing PTBs (green fluorescence). The long arrows represent the PTBs with non-GFP-Rnd3. ROS levels were significantly reduced in PE-PTBs following GFP-Rnd3 overexpression as determined by DHE staining. NS indicates non-specific staining. The scale bar represents 25 *μ*m. (**B**) 8-isoprostane level detection in the C-PTBs and P-PTBs following treatment of Ad-GFP or Ad-GFP-Rnd3. (**C**) Ad-GFP-hRnd3 improved *UCP2* mRNA transcripts in the PTBs.**(D)** Rnd3 overexpression rescued PPARγ-UCP2 signaling in PE-PTBs. Mitochondrial dysfunction in PE-PTBs was partially rescued by Ad-GFP-hRnd3, as determined by the increases in proton leak associated OCR (**E**) and respiratory control ratio (**F**). C-PTBs indicates control-primary trophoblasts; P-PTBs, PE-primary trophoblasts.

### Downregulation of the expression levels of PPARγ and UCP2 were observed in human placentae with PE

The expression levels of PPARγ and UCP2 were evaluated in human PE placental tissues. The experiments aimed to offer additional insight into the clinical relevance of regulating the PPARγ/UCP2 pathway by Rnd3 in PE. PPARγ protein levels were downregulated in PE placentae. Consistent with the post-transcriptional regulation of PPARγ by Rnd3, the corresponding mRNA levels of *PPARγ* indicated no significant changes between the control and PE groups (Fig 9A-B). The mRNA transcripts of *UCP2* and its protein levels were reduced in human PE placentae (Fig 9A and C).

**Figure 9.**
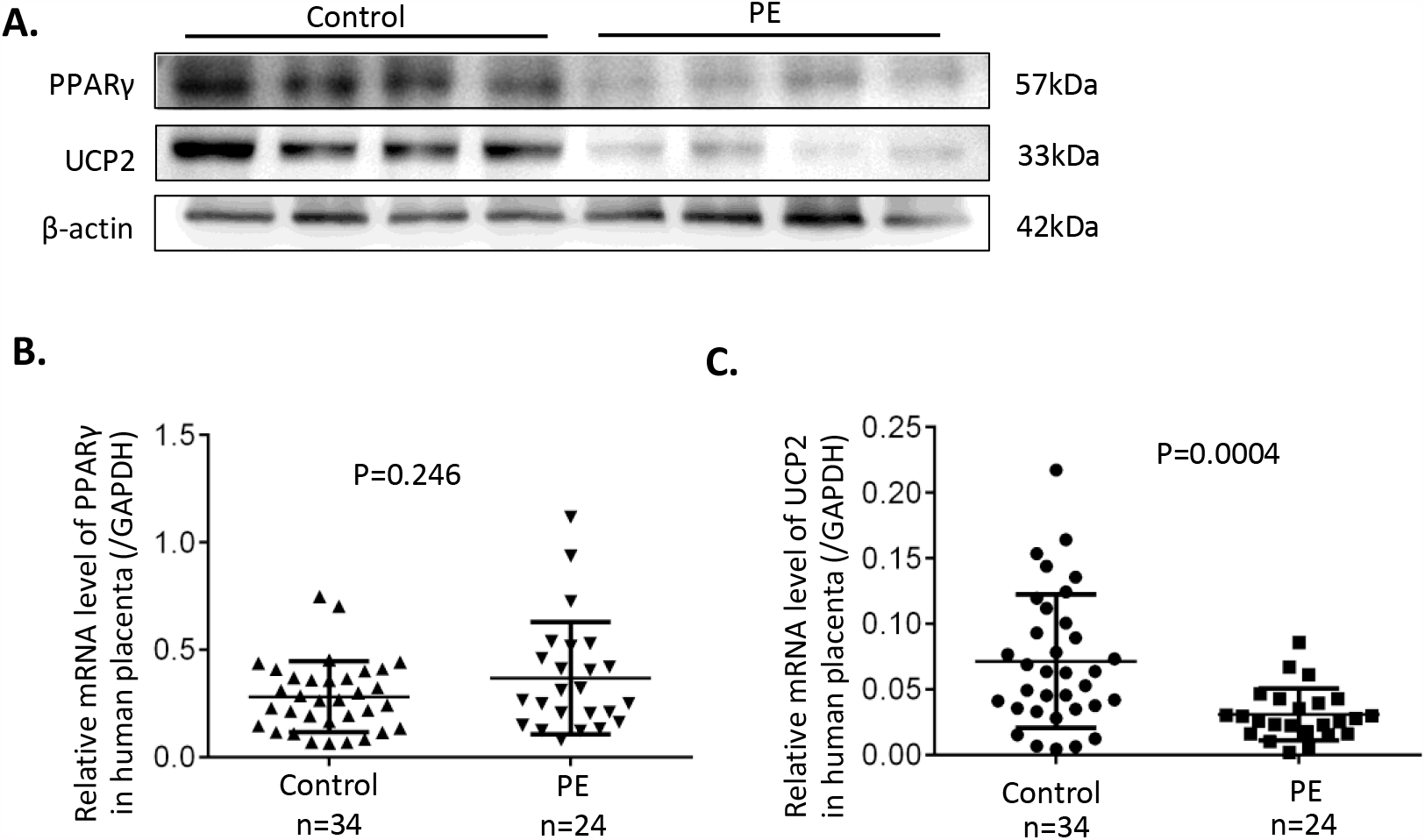
The downregulation of the PPARγ-UCP2 cascade is confirmed in human placentae with PE. **(A)** Representative protein expression levels of PPARγ and UCP2 as determined by western blot analysis. The levels of the two proteins were downregulated in the placentas with PE. (**B**) Relative mRNA levels of *PPARγ* from 34 healthy placentae and 24 placentae with PE. (**C**) Relative mRNA levels of *UCP2* from 34 healthy placentae and 24 placentae with PE.

## Discussion

PE has been suggested as a mitochondrial disorder since the late 1980s(32). Proteome analysis of PE placentae revealed that the abnormal expression of the respiratory complex proteins was associated with ROS’s excessive generation in the mitochondrial respiratory chain(33). Given that mitochondrial ROS is an important contributor to oxidative stress and inflammation, the reason causing placental mitochondrial ROS is critical and remains to be revealed. This study uncovers a novel understanding of placental oxidative stress in PE that Rnd3-mediated PPARγ/UCP2 cascade participates in mitochondrial dysfunction regulation.

In PE’s pathological condition, Rnd3 promotes trophoblasts’ proliferation and migration under regulations by lncRNAs HOXA11-AS and TUG1(27, 28). In contrast to these observations, Rnd3 has also been proposed as a stimulator of trophoblast cell invasion in PE, under the repression caused by miR-182-5p(29). Our study confirmed the regulatory role of Rnd3 in PE and revealed the novel function of Rnd3 in maintaining the mitochondrial respiratory chain. The supply of Rnd3 in PE patient-derived primary trophoblasts significantly improved mitochondrial function and repressed oxidative stress (Fig 8), suggesting the potential role of Rnd3 as a pharmacological target for PE.

Reductions in *Rnd3* mRNA and protein levels were observed in human PE placentas. One possible factor regulating Rnd3 expression could be hypoxia. In BeWo cells, the expression of Rnd3 was stimulated by cAMP/PKA signaling, however, was inhibited under hypoxic condition(34). These suggest that, in PE pathogenesis, failed spiral artery remodeling causes placental ischemia and hypoxia, and thereby contributing to the downregulation of Rnd3 in placenta. Moreover, IGFBP1 was found downregulated in the serum of women with PE, and the downregulation of IGFBP1 may participated in pathogenesis of PE by hindering trophoblast migration through ROCK1 and ROCK2 pathway(35). Considering that ROCK1 phosphorylates and stabilizes RND3 protein, these may explain the reduce of Rnd3 in PE patients(36).

Sun’s group reported increases in the mRNA levels of *Rnd3* in the placentas from PE patients, which is opposite to our data(27, 28). To verify the expression pattern of Rnd3 in PE, we detected both mRNA and protein expressions of Rnd3, and confirmed the downregulation of Rnd3 in the PE placentas. One reason of the controversy could be that the specimens were obtained at different sites of placentas. Sun’s group collected placental tissue samples from the central area of the placenta’s maternal surface(27). In our study, the placental tissues were collected from the mid-section between the chorionic and maternal basal surfaces at four different positions 2 cm from central umbilical cord. The distribution of villous structure within placenta is variable. Cytotrophoblasts and syncytiotrophoblasts cover the surface of villi, and therefore the tissues from placenta’s maternal surface consist of large amount of trophoblasts. Whereas the tissues between the chorionic and maternal surfaces were more likely consist of villous core stroma, including mesenchymal cells, hofbauer cells and fibroblasts, as well as trophoblastic layers. Meanwhile, blood vessels were also non-uniformly distributed, with main branches enriched in the central area of placenta. Therefore, the protein expressions from different sites of placentas were highly variable. Besides, samples were obtained at different gestation weeks (33.93 ± 3.231, n=60 vs. 36.1 ± 2.8, n=24; p=0.005), which may also contribute to the difference in Rnd3 expressions.

UCP2 exerts protective properties against oxidative damage by reducing ROS generation via decreasing mitochondrial proton gradient and local oxygen availability(37). PPARγ and its ligands directly activate the *UCP2* gene promoter via the E-box element(31). By recovering the mitochondrial membrane potential and reducing ROS generation, PPARγ improves trophoblast cells’ survival under hypoxic conditions (38). In the present study, reduced PPARγ/UCP2 signaling was detected in human PE placentas, contributing to mitochondrial defect and oxidative stress in PE primary trophoblasts. However, excessive stimulation of PPARγ/UCP2 has also been implicated in pathological placenta with maternal nutrient restriction by enhancing fatty acid metabolism and limiting glucose utilization(39). Therefore, the homeostasis of the PPARγ/UCP2 axis is critical in maintaining normal placental function.

It is interesting to note that Rnd3 facilitates the protein accumulation of PPARγ without causing the change of PPARγ mRNA transcripts. Therefore, immunoprecipitation was performed to investigate the underlying mechanism, and the direct interaction of protein molecules of RND3 and PPARγ was observed. Meanwhile, the co-localization of the two proteins was visualized in nuclei and cytoplasm (Fig 6G). Given that the dynamics of PPARγ depends on ligand-induced transcriptional activation and ubiquitin-proteasome dependent degradation, it is reasonable that Rnd3 may stabilize PPARγ in a post-translational manner.

The present study has several limitations. First, even though we have used two cell models, including PE primary cell and the transformed BeWo cell line, to mimic the Rnd3 downregulation in PE patients’ placentas. It is not precisely clear whether Rnd3 deficiency could cause PE *in vivo*. Future studies using the *Rnd3* gene knockout animal model will provide stronger supports for the role of Rnd3 in PE pathogenesis. Second, even though we have proved the direct interaction of RND3 and PPARγ protein molecules; however, the more precise post-translational regulation mechanism also needs further investigation.

## Conclusions

We identified here the downregulation of Rnd3 in placental trophoblasts in patients with PE and proved that Rnd3 deficiency could cause mitochondrial defects and oxidative stress in trophoblasts. Supply of Rnd3 in PE primary trophoblasts attenuated mitochondrial dysfunction and oxidative stress. The possible underlying mechanism is that Rnd3 interacts with PPARγ and stabilizes PPARγ protein, causing stimulation of PPARγ/UCP2 cascade. In conclusion, our study indicates the novel role of Rnd3, providing new insights into PE pathogenesis.

## Abbreviations

ACOG: American College of Obstetricians and Gynecologists
CK7: cytokeratin 7
CTB: cytotrophoblast
DHE: dihydroethidium
FBS: fetal bovine serum
IUGR: fetal intrauterine growth restriction
LOC: loss of cristae
NRF1: nuclear respiratory factor 1
OCR: oxygen consumption rates
PE: preeclampsia
PGC1-α: PPARγ coactivator 1-α
PPARα: peroxisome proliferator-activated receptor α
PPARγ: peroxisome proliferators-activated receptor γ
PTBs: primary trophoblasts
ROS: reactive oxygen species
SOD1: superoxide dismutase 1
SOD2: superoxide dismutase 2
STB: syncytiotrophoblast
TFAM: mitochondrial transcription factor A
TUNEL: terminal deoxynucleotidyl transferase dUTP nick end labeling
UCP2: mitochondrial uncoupling protein 2

## Disclosure of interests

None.

## Contribution to authorship

X.Y. designed the experiments; L.H., Y.M. and X.Y. conducted most experiments; L.C. collected data in Fig. 9; Y.S., X.C., F.S., L.X., J.C., Y.L., C.Y. and X.Y. analyzed the data; X.Y. wrote the manuscript; J.C., M.Z., Z.W. and X.Y. revised the manuscript. All authors contributed to the final manuscript.

## Funding

This study was supported by the following funding sources: the Major Science and Technology Program of Hainan Province (ZDKJ2017007), the National Natural Science Foundation of China (81771609, 81601317, 81960283, and 81971415), the Natural Science Foundation of Guangdong Province (2017A030313584, 2019A1515010290 and 2019A1515010019), the Special Fund for Cooperative Innovation and Platform Environment Construction (2015B050501006), the Outstanding Youth Development Scheme of Nanfang Hospital Southern Medical University (2018J010).

---

**Table 1. Clinical characteristics of the PE group and of the healthy control group**. Clinical data of blood pressure, BMI and proteinuria was collected at the time of PE diagnosis.

**Figure S1.**
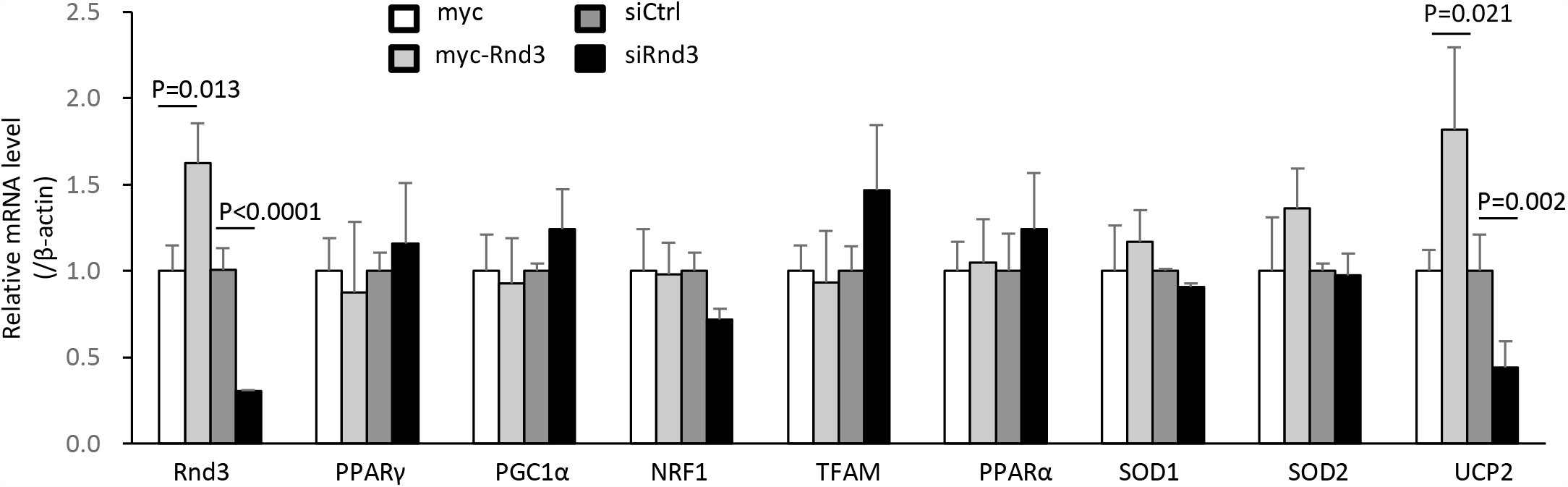
Relative mRNA expression levels of the mitochondrion-related genes in BeWo cells. Rnd3 promoted *UCP2* mRNA level. Manipulation of Rnd3 expression resulted in no change in the mRNA levels of *PPARγ, PGC1α, NRF1, TFAM, PPARα, SOD1* and *SOD2*.

**Figure S2.**
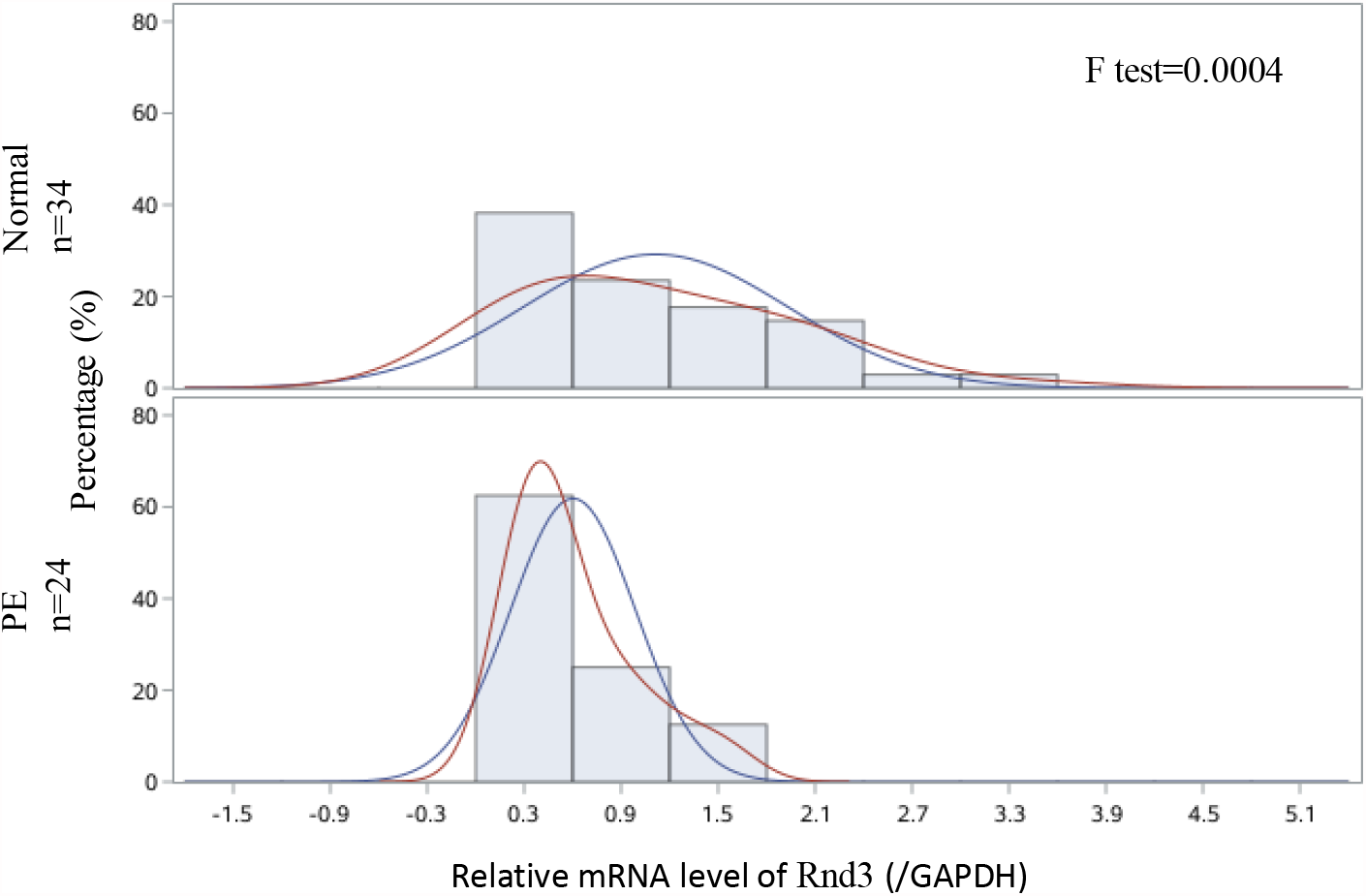
Tests of homogeneity of variances for relative mRNA levels of *Rnd3* in human placentas.

**Figure S3.**
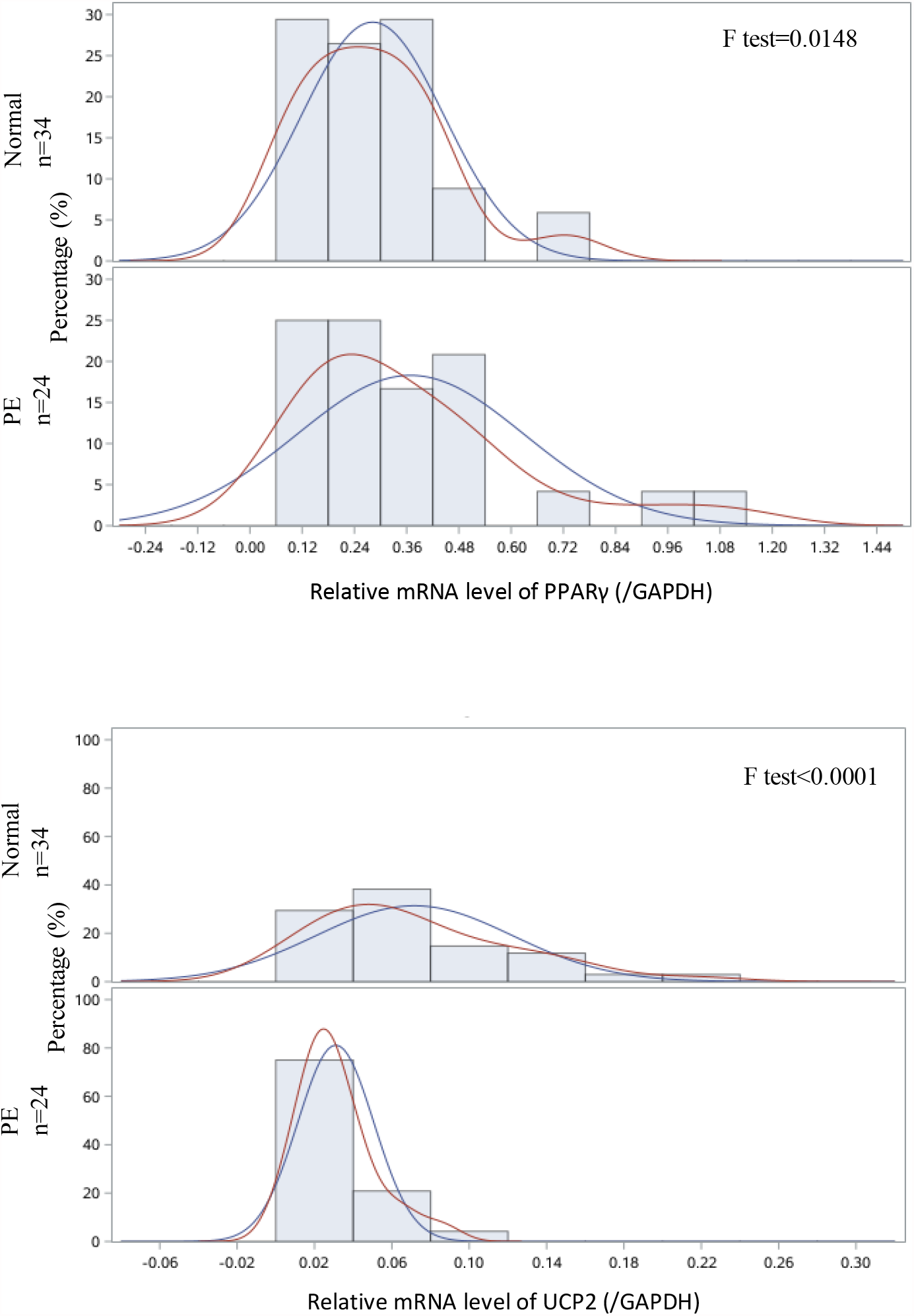
**Tests of homogeneity of variances for relative mRNA levels of *PPARγ* and *UCP2* in human placentas.**

## Notes

### Competing Interest Statement

The authors have declared no competing interest.

## References

1. Rana, S., Lemoine, E., Granger, J. P., and Karumanchi, S. A. (2019) Preeclampsia: Pathophysiology, Challenges, and Perspectives. Circ Res 124, 1094–1112

2. Steegers, E. A., von Dadelszen, P., Duvekot, J. J., and Pijnenborg, R. (2010) Pre-eclampsia. Lancet 376, 631–644

3. Hladunewich, M., Karumanchi, S. A., and Lafayette, R. (2007) Pathophysiology of the clinical manifestations of preeclampsia. Clin J Am Soc Nephrol 2, 543–549

4. Birben, E., Sahiner, U. M., Sackesen, C., Erzurum, S., and Kalayci, O. (2012) Oxidative stress and antioxidant defense. World Allergy OrganJ 5, 9–19

5. San Juan-Reyes, S., Gomez-Olivan, L. M., Islas-Flores, H., and Dublan-Garcia, O. (2020) Oxidative stress in pregnancy complicated by preeclampsia. Arch Biochem Biophys 681, 108255

6. Myatt, L. (2010) Review: Reactive oxygen and nitrogen species and functional adaptation of the placenta. Placenta 31 Suppl, S66–69

7. Sanchez-Aranguren, L. C., Prada, C. E., Riano-Medina, C. E., and Lopez, M. (2014) Endothelial dysfunction and preeclampsia: role of oxidative stress. Front Physiol 5, 372

8. Wang, Y., and Walsh, S. W. (1998) Placental mitochondria as a source of oxidative stress in pre-eclampsia. Placenta 19, 581–586

9. D’Souza, V., Rani, A., Patil, V., Pisal, H., Randhir, K., Mehendale, S., Wagh, G., Gupte, S., and Joshi, S. (2016) Increased oxidative stress from early pregnancy in women who develop preeclampsia. Clin Exp Hypertens 38, 225–232

10. Vishnyakova, P. A., Volodina, M. A., Tarasova, N. V., Marey, M. V., Tsvirkun, D. V., Vavina, O. V., Khodzhaeva, Z. S., Kan, N. E., Menon, R., Vysokikh, M. Y., and Sukhikh, G. T. (2016) Mitochondrial role in adaptive response to stress conditions in preeclampsia. Sci Rep 6, 32410

11. Salgado, S. S., and Salgado, M. K. R. (2011) Structural changes in pre-eclamptic and eclamptic placentas--an ultrastructural study. J Coll Physicians Surg Pak 21, 482–486

12. Janani, C., and Ranjitha Kumari, B. D. (2015) PPAR gamma gene--a review. Diabetes Metab Syndr 9, 46–50

13. Corona, J. C., and Duchen, M. R. (2016) PPARgamma as a therapeutic target to rescue mitochondrial function in neurological disease. Free Radic Biol Med 100, 153–163

14. Waite, L. L., Louie, R. E., and Taylor, R. N. (2005) Circulating activators of peroxisome proliferator-activated receptors are reduced in preeclamptic pregnancy. J Clin Endocrinol Metab 90, 620–626

15. Barak, Y., Nelson, M. C., Ong, E. S., Jones, Y. Z., Ruiz-Lozano, P., Chien, K. R., Koder, A., and Evans, R. M. (1999) PPAR gamma is required for placental, cardiac, and adipose tissue development. MolCell 4, 585–595

16. Barak, Y., Liao, D., He, W., Ong, E. S., Nelson, M. C., Olefsky, J. M., Boland, R., and Evans, R. M. (2002) Effects of peroxisome proliferator-activated receptor delta on placentation, adiposity, and colorectal cancer. ProcNatlAcadSciUSA 99, 303–308

17. Schaiff, W. T., Carlson, M. G., Smith, S. D., Levy, R., Nelson, D. M., and Sadovsky, Y. (2000) Peroxisome proliferator-activated receptor-gamma modulates differentiation of human trophoblast in a ligand-specific manner. J Clin Endocrinol Metab 85, 3874–3881

18. Waite, L. L., Person, E. C., Zhou, Y., Lim, K. H., Scanlan, T. S., and Taylor, R. N. (2000) Placental peroxisome proliferator-activated receptor-gamma is up-regulated by pregnancy serum. J Clin Endocrinol Metab 85, 3808–3814

19. McCarthy, F. P., Drewlo, S., Kingdom, J., Johns, E. J., Walsh, S. K., and Kenny, L. C. (2011) Peroxisome proliferator-activated receptor-gamma as a potential therapeutic target in the treatment of preeclampsia. Hypertension 58, 280–286

20. Liu, B., Lin, X., Yang, X., Dong, H., Yue, X., Andrade, K. C., Guo, Z., Yang, J., Wu, L., Zhu, X., Zhang, S., Tian, D., Wang, J., Cai, Q., Chen, Q., Mao, S., Chen, Q., and Chang, J. (2015) Downregulation of RND3/RhoE in glioblastoma patients promotes tumorigenesis through augmentation of notch transcriptional complex activity. Cancer Med 4, 1404–1416

21. Yue, X., Yang, X., Lin, X., Yang, T., Yi, X., Dai, Y., Guo, J., Li, T., Shi, J., Wei, L., Fan, G. C., Chen, C., and Chang, J. (2014) Rnd3 haploinsufficient mice are predisposed to hemodynamic stress and develop apoptotic cardiomyopathy with heart failure. Cell DeathDis 5, e1284

22. Yue, X., Lin, X., Yang, T., Yang, X., Yi, X., Jiang, X., Li, X., Li, T., Guo, J., Dai, Y., Shi, J., Wei, L., Youker, K. A., Torre-Amione, G., Yu, Y., Andrade, K. C., and Chang, J. (2016) Rnd3/RhoE Modulates Hypoxia-Inducible Factor 1alpha/Vascular Endothelial Growth Factor Signaling by Stabilizing Hypoxia-Inducible Factor 1alpha and Regulates Responsive Cardiac Angiogenesis. Hypertension 67, 597–605

23. Yang, X., Wang, T., Lin, X., Yue, X., Wang, Q., Wang, G., Fu, Q., Ai, X., Chiang, D. Y., Miyake, C. Y., Wehrens, X. H. T., and Chang, J. (2015) Genetic deletion of Rnd3/RhoE results in mouse heart calcium leakage through upregulation of protein kinase A signaling. Circ Res 116, e1–e10

24. Dai, Y., Song, J., Li, W., Yang, T., Yue, X., Lin, X., Yang, X., Luo, W., Guo, J., Wang, X., Lai, S., Andrade, K. C., and Chang, J. (2019) RhoE Fine-Tunes Inflammatory Response in Myocardial Infarction. Circulation 139, 1185–1198

25. Lin, X., Liu, B., Yang, X., Yue, X., Diao, L., Wang, J., and Chang, J. (2013) Genetic deletion of Rnd3 results in aqueductal stenosis leading to hydrocephalus through up-regulation of Notch signaling. Proc Natl AcadSci USA 110, 8236–8241

26. Moses, E., Johnson, M., East, C., Dyer, T., Roten, L., Proffitt, J., Fenstad, M., Aalto-Viljakainen, T., Makikallio, K., Heinonen, S., Kajantie, E., Kere, J., Laivuori, H., Austgulen, R., Blangero, J., and Brennecke, S. (2012) OS077. The chromosome 2q22 preeclampsia susceptibility locus reveals shared novel risk factors for CVD. Pregnancy Hypertens 2, 219–220

27. Xu, Y., Wu, D., Liu, J., Huang, S., Zuo, Q., Xia, X., Jiang, Y., Wang, S., Chen, Y., Wang, T., and Sun, L. (2018) Downregulated lncRNA HOXA11-AS Affects Trophoblast Cell Proliferation and Migration by Regulating RND3 and HOXA7 Expression in PE. Mol Ther Nucleic Acids 12, 195–206

28. Xu, Y., Ge, Z., Zhang, E., Zuo, Q., Huang, S., Yang, N., Wu, D., Zhang, Y., Chen, Y., Xu, H., Huang, H., Jiang, Z., and Sun, L. (2017) The lncRNA TUG1 modulates proliferation in trophoblast cells via epigenetic suppression of RND3. Cell Death Dis 8, e3104

29. Fang, Y. N., Huang, Z. L., Li, H., Tan, W. B., Zhang, Q. G., Wang, L., and Wu, J. L. (2018) Highly expressed miR-182-5p can promote preeclampsia progression by degrading RND3 and inhibiting HTR-8/SVneo cell invasion. Eur Rev Med Pharmacol Sci 22, 6583–6590

30. Yue, X., Sun, Y., Zhong, M., Ma, Y., Wei, Y., Sun, F., Xiao, L., Liu, M., Chen, J., Lai, Y., Yan, C., Huang, L., and Yu, Y. (2018) Decreased expression of fibroblast growth factor 13 in early-onset preeclampsia is associated with the increased trophoblast permeability. Placenta 62, 43–49

31. Medvedev, A. V., Snedden, S. K., Raimbault, S., Ricquier, D., and Collins, S. (2001) Transcriptional regulation of the mouse uncoupling protein-2 gene. Double E-box motif is required for peroxisome proliferator-activated receptor-gamma-dependent activation. J BiolChem 276, 10817–10823

32. Torbergsen, T., Oian, P., Mathiesen, E., and Borud, O. (1989) Pre-eclampsia--a mitochondrial disease? Acta Obstet Gynecol Scand 68, 145–148

33. Shi, Z., Long, W., Zhao, C., Guo, X., Shen, R., and Ding, H. (2013) Comparative proteomics analysis suggests that placental mitochondria are involved in the development of pre-eclampsia. PLoS One 8, e64351

34. Collett, G. P., Goh, X. F., Linton, E. A., Redman, C. W. G., Sargent, I. L. (2012) RhoE is regulated by cyclic AMP and promotes fusion of human BeWo choriocarcinoma cells. PLoS One 7, e30453

35. Saso, J., Shields, S., Zuo, Y., Chakraborty, C. (2012) Role of Rho GTPases in human trophoblast migration induced by IGFBP1. Biol Reprod 86, 1–9.

36. Jie, W., Andrade, K. C., Lin, X., Yang, X., Yue, X., Chang, J. (2015) Pathophysiological Functions of Rnd3/RhoE. Compr Physiol 6, 169–186.

37. Cadenas, S. (2018) Mitochondrial uncoupling, ROS generation and cardioprotection. Biochim Biophys Acta Bioenerg 1859, 940–950

38. Kohan-Ghadr, H. R., Kilburn, B. A., Kadam, L., Johnson, E., Kolb, B. L., Rodriguez-Kovacs, J., Hertz, M., Armant, D. R., and Drewlo, S. (2019) Rosiglitazone augments antioxidant response in the human trophoblast and prevents apoptosisdagger. Biol Reprod 100, 479–494

39. Yiallourides, M., Sebert, S. P., Wilson, V., Sharkey, D., Rhind, S. M., Symonds, M. E., and Budge, H. (2009) The differential effects of the timing of maternal nutrient restriction in the ovine placenta on glucocorticoid sensitivity, uncoupling protein 2, peroxisome proliferator-activated receptor-gamma and cell proliferation. Reproduction 138, 601–608

